# Host exposure to symbionts and ecological drift generate divergence in parasite community assembly

**DOI:** 10.1101/2021.11.19.469294

**Authors:** Rita L. Grunberg, Brooklynn N. Joyner, Charles E. Mitchell

## Abstract

The initial colonization of a host by symbionts, ranging from parasites to mutualists, can generate priority effects that alter within-host interactions and the trajectory of parasite community assembly. At the same time, variation in parasite communities among hosts can also stem from stochastic processes. Community ecology theory posits that multiple processes (e.g. dispersal, selection and drift) interact to generate variation in community structure, but these processes are rarely considered simultaneously during community assembly. To test the role of these processes in a parasite community, we experimentally simulated dispersal of three symbionts by factorially inoculating individual plants of tall fescue with two foliar fungal parasites, *Colletotrichum cereale* and *Rhizoctonia solani*, and a hypothesized mutualist endophyte, *Epichloë coenophiala*. We then tracked parasite infections longitudinally in the field. After the initial inoculations, hosts were exposed to a common pool of parasites in the field, which we expected to cause parasite communities to converge towards a similar community state. To test for convergence, we analyzed individual hosts’ parasite community trajectories in multivariate space. In contrast to our expectation, there was no signal of convergence. Instead, parasite community trajectories generally diverged over time between treatment groups and the magnitude of divergence depended on the symbiont species inoculated. Parasite communities of hosts that were inoculated with only the mutualist, *Epichloë*, showed significant trends of divergence relative to all other symbiont inoculation treatments. In contrast, hosts inoculated with only *Rhizoctonia* did not exhibit clear trends of divergence when compared to other parasite inoculations. Further, co-inoculation with both parasite species resulted in faster rates of divergence and greater temporal change in parasite communities relative to hosts inoculated with only the parasite *Colletotrichum*. As predicted by existing theory, parasite communities showed evidence of drift during the beginning of the experiment, which contributed to among-host divergence in parasite community structure. Overall, these data provide evidence that initial dispersal of symbionts produced persistent changes in parasite community structure via ecological selection, that drift was important during the early stages of parasite community assembly, and together, dispersal, selection and drift resulted in parasite community divergence.

**Open Research statement:** The data and code that support the findings of this study are available through Zenodo at https://doi.org/10.5281/zenodo.5714452

## Introduction

Most parasite infections occur within the context of a diverse community of parasites. Coinfections, i.e., simultaneous infections by multiple parasite species or strains, can drive disease outcomes within and among host individuals (Hoverman et al. 2013, Halliday et al. 2017, Marchetto and Power 2018, Clay et al. 2019). Still, parasite communities are spatially and temporally diverse, and it is less clear as to what degree coinfections contribute to this variation in parasite community structure among host individuals. Variation in community structure has been attributed to four processes: dispersal, selection, drift, and speciation/diversification (Vellend 2010). Within a regional pool of parasites, dispersal rates differ considerably among species, generating variation in the timing of infections among hosts. Further, the successful establishment of colonizing parasite species is contingent on niche-based processes, which consists of environmental filtering and species interactions (i.e., collectively, selection). Parasite community structure is also driven by stochastic processes (i.e., birth/death and reproduction), but less is known about the importance of drift (Seabloom et al. 2015). Finally, speciation is likely less important during the lifespan of an individual host infected with macroparasites. Ultimately, these four processes will interact to generate variation in parasite community structure among hosts.

Initial variation in species composition among habitats can be generated by differences in species dispersal with consequences for selection (Shurin 2001). The outcome of species interactions is often contingent on the order of species arrival and this may generate priority effects with long-lasting effects on community structure (Fukami 2015). Within parasite communities, priority effects are likely to occur between parasite species that have a high degree of niche overlap and modify the host environment/resources in ways that facilitate or impede infections by other species (Vannette and Fukami 2014). For example, the first parasite to colonize a host can stimulate the host immune system in ways that alters the host’s susceptibility to other parasite infections (Graham 2008, Syller and Grupa 2016) and consequently alters the overall strength of species interactions (Hoverman et al. 2013, Halliday et al. 2018). Evidence for priority effects have been demonstrated in a diverse array of host-parasite systems (Benesh and Kalbe 2016, Halliday et al. 2017, 2020, Zilio and Koella 2020), and can have consequences for the distribution (Halliday et al. 2017) and abundance (Wuerthner et al. 2017) of parasitic species. However, priority effects do not always have lasting effects on community structure (Fukami 2015), making it important to consider to what degree a host’s initial exposure to parasites results in persistent changes in parasite community structure during community assembly.

When evaluating the effects of initial parasite infections on parasite community structure it is also necessary to consider the effects of other symbiont species (i.e., host associated microbes, which include parasites, commensals, and mutualists) infecting a host. This is because parasites commonly interact with a diverse community of symbionts that have the potential to modify parasite transmission and infection intensity (Hopkins et al. 2017). Specifically, mutualistic symbionts are ubiquitous across hosts and can confer disease resistance by inhibiting parasite growth (Kruger 2020) and reproduction (Fernandez et al. 1991, Saikkonen et al. 2010). Some mutualists are known to stimulate immune pathways associated with defense against certain parasites (Saikkonen et al. 2013, Bastias et al. 2017, Pérez et al. 2020). However, the importance of mutualists in mediating parasite community structure remains less clear because their overall effects on host susceptibility or parasite growth rates are hard to predict across parasite species within a community. For example, some subsets of parasite species may respond directly to the specific immune pathways elicited by the mutualist, but other parasite species may respond indirectly when mutualists modify within host interactions (Adame-Álvarez et al. 2014, O’Keeffe et al. 2021). In the end, the outcome of within host interactions may be contingent on the presence of other symbionts. Thus, the general trajectory of parasite community assembly has the potential to change when in coinfection with mutualists.

Although the role of species interactions in shaping parasite diversity has been explored, relatively little is known about the importance of drift in generating variation in parasite communities (Seabloom et al. 2015). Nonetheless, drift is an important driver of community dissimilarity patterns across free-living systems (Germain et al. 2013, Gilbert and Levine 2017), and considering its effects, along with deterministic processes, is critical for community ecology (Vellend 2010). All else being equal, stochastic fluctuations of the relative abundance of certain parasitic taxa will occur in naturally infected hosts and this will impact community structure when population sizes are small (Vellend 2010), suggesting that drift is important during the initial stages of infection (Seabloom et al. 2015). Consequently, it is important to consider how among host variation in parasite community structure related to drift can influence parasite diversity patterns and how this relationship may change as parasite communities develop.

To test for the effects of host exposure history to symbionts and drift on parasite diversity, we aim to determine 1) whether the order of arrival of parasite species (i.e., variation in dispersal and selection) alters parasite community patterns, 2) how community patterns change when in coinfection with a mutualist endophyte (i.e., selection), and 3) the role of drift in generating variation in parasite communities among host individuals. To quantify changes in parasite community patterns, we inoculated individual plants of tall fescue (*Lolium arundinaceum*) in the lab with a factorial combination of three fungal symbionts (2 parasites, 1 mutualistic endophyte), deployed the hosts in the field, then conducted longitudinal disease surveys in the field. Because hosts were exposed to transmission from a common pool of parasites in the field, the parasite communities might be expected to ultimately converge towards a similar community state despite the experimentally imposed differences in initial communities. Thus, our goal was to evaluate whether effects from an initial inoculation event from a diverse array of symbionts have long-lasting effects on parasite communities within individual hosts.

## Methods

### Study system

We inoculated individual plants of tall fescue, *L. arundinaceum*, with a factorial combination of three symbionts: two foliar fungal parasites, *Colletotrichum cereale* and *Rhizoctonia solani*, and a fungal endophyte, *Epichloë coenophiala*. The parasite *Colletotrichum cereale* is a causative agent of the disease known as anthracnose, and is hemibiotrophic, meaning it initially extracts resources from living host cells (i.e., biotrophic) and later switches to killing host cells and extracting resources from the dead tissue (i.e., necrotrophic). Transmission of this parasite occurs chiefly through rain splash, which facilitates the dispersal of its mucilaginous spores. The fungal parasite *Rhizoctonia solani* is a facultative necrotroph; it can survive within the soil and infects and extracts resources from host plants as a necrotroph. *Rhizoctonia* is transmitted primarily by hyphae and sclerotia. The third symbiont manipulated in our experiment, *Epichloë coenophiala*, is a systemic fungal endophyte that is transmitted vertically by seed. Generally, *Epichloë* is considered a mutualist of tall fescue with some defensive properties against herbivory (Clay 1988, Saikkonen et al. 2010). But, ultimately its interactions with its host may be context specific with respect to disease, for example, *Epichloë* is proposed to facilitate biotrophs and inhibit necrotrophs (Saikkonen et al. 2013).

### Experiment

In total, we factorially inoculated 136 plants with three symbionts: *Colletotrichum cereale* (parasite), *Rhizoctonia solani* (parasite), and *Epichloë coenophialia* (mutualist), which yielded eight treatment groups (sample sizes summarized in Table S1). *Epichloë* is transmitted from infected seed, so we used *Epichloë-free* and *Epichloë-infected* seeds of the cultivar KY-31. Then we grew plants from seed under greenhouse conditions for eight weeks and exposed plants to the parasite inoculum treatments. For the *R*. *solani* inoculation, we placed a mycelium PDA plug at the base of the oldest living leaf, following O’Keeffe et al. 2021. The *C. cereale* inoculations were implemented according to Beirn et al., 2015, where we applied a spore solution on the oldest living leaf. Further details on the symbiont inoculations procedures are described in Appendix I. After the inoculation procedures, we moved plants into the field and maintained plants in pots placed into individual holes within a 1.5 x 12 m plot located at Widener Farm in the Duke Forest Teaching and Research Laboratory in Orange Co., North Carolina, USA (Appendix II). Plants were placed into holes randomly within the plot and each plant was relocated to a new hole every week to homogenize parasite exposure.

To quantify the parasite community within each plant, we surveyed all plants visually for disease symptoms (lesions and pustules) twice per week for six weeks in the field. Disease symptoms are an epidemiologically relevant measure of parasite infection because, transmission of our parasites from an infected leaf to another leaf requires production of a lesion or pustule, as these are the source of parasite propagules. Disease surveys occurred from 19-Sept-2019 to 31-Oct-2019, corresponding to the seasonal period which the three common parasite species are present in the resident host population (Halliday et al. 2017). Throughout these surveys, parasite infections were characterized at the leaf level since infections are localized within an individual leaf (Halliday et al 2017). The lifespan of inoculated leaves varied among plants in the field (some inoculated leaves died after the first disease survey, Figure S1), and their relative contribution to an individual plant’s parasite severity measure ranged from 2% to 35%.

We recorded parasite infection severity (i.e., % area of leaf damaged by a parasite species) on all leaves in a tiller (i.e., individual grass shoot originating from a plant) within a plant. If a tiller died or produced a daughter tiller, a new tiller was selected. All leaves in a tiller were surveyed longitudinally for parasites. The relative age of each leaf in a tiller was also monitored: pre-existing leaves prior to transplant into the field were assigned to age 0 at the initial survey event, and each newly emerged leaf during the survey was assigned to age 0. We identified previously surveyed leaves based on their vertical order within a plant.

During the disease surveys, we recorded the presence of four foliar fungal parasite species. These parasites were distinguished from each other based on visual disease symptoms that align with causal parasite species in this system (Halliday et al. 2017, 2018). The parasites and their associated symptoms observed include: *Colletotrichum cereale* which chiefly although not exclusively causes the disease anthracnose, *Rhizoctonia solani* causes brown patch, *Puccinia coronata* causes crown rust, and *Pyricularia grisea* causes gray leaf spot. Hereafter, rather than the disease symptoms, we refer to these causal parasite species.

On the final disease survey date, we harvested plants for above-ground biomass and to confirm *Epichloë* infection status via immunoblot assay (Agrinostics Ltd. Co, Watkinsville, GA, USA). Overall, 5% to 48% of the plants from the *Epichloë* treatment groups tested positive for *Epichloë* infection (Table S1). Notably, within our field site, tall fescue is frequently infected with *Epichloë* and 85% (628 total plants assayed in another experiment) of the resident plants tested positive for the endophyte in a 2019 field study (B. N. Joyner, unpublished data).

### Analysis

The aim of our study was to quantify changes in parasite community structure in response to the host’s initial symbiont inoculation. To accomplish this, we took a multivariate approach and focused on patterns of dissimilarity among parasite communities within plants and analyzed community trajectories following the framework of De Cáceres et al. 2019. We established patterns of community dissimilarity by generating ordinations using non-metric multidimensional scaling (NMDS) on Bray-Curtis distances based on parasite infection severity at the plant-level (i.e., parasite infection severity (% area of leaf damaged) averaged across all leaves nested within a plant) in the ‘vegan’ R package. Although these infections are localized within a leaf (Halliday et al. 2018) and inoculations were conducted on a single leaf, we used plant-level parasite severity because infections of leaves within a plant are not statistically independent and inoculation treatments were randomized and implemented at the plant level.

When calculating plant-level infection severity for the parasite trajectory analyses, we excluded 454 leaves that never showed parasite disease symptoms during the field survey (n = 788 total leaves monitored for infection) because these leaves had no parasite community trajectory to incorporate, so in total we used longitudinal infection measurements from 334 infected leaves for our analyses. Excluding leaves that were never infected from our analysis did not alter our main results. We also excluded three plants that never showed disease symptoms from all analyses (Table S1).

On the first disease survey, all hosts showed no visual disease symptoms, which was expected, so to retain these “no parasite community” observations in these analyses we included a dummy variable in the parasite community matrix. The inclusion of the dummy variable causes hosts with no observed parasite community to cluster together, so all hosts started at the same community state and thus had the same NMDS coordinates. The parasite matrix contained 4 fungal parasite species and the dummy variable. Because parasite infection severity varied considerably across parasite species (e.g., maximum observed infection severity was 90% of a leaf’s area infected by *Rhizoctonia*, 50% for *Colletotrichum*, 3% for *Pyricularia*, and 2% for *Puccinia*), we standardized infection severity based on each species maximum to facilitate a more uniform weighting across parasite species.

### Multivariate patterns of parasite community structure

We first evaluated whether parasite community patterns differed in response to symbiont inoculation treatments and across survey days using a PERMANOVA (function ‘Adonis’) based on Bray-Curtis distance on the entire dataset. In the PERMANOVA, we included symbiont inoculation treatment and survey day in our model, and permutations in the PERMANOVA were constrained within an individual plant to account for repeated sampling. After the group wide PERMANOVA, we evaluated differences between all symbiont inoculation treatment groups using pairwise PERMANOVAs and these p-values were adjusted for multiple comparisons using Benjamini-Hochberg corrections. We also tested for multivariate homogeneity of dispersion among inoculation treatments and survey days using the ‘betadisper’ function and an ANOVA. To further assess whether dispersion varied between inoculation treatments we used Tukey HSD post hoc comparisons.

### Temporal change in parasite community structure

By potentially altering parasite growth, colonization, and extinction rates, due to selection, the identity of symbionts that initially infect a host may also affect how much its parasite community subsequently changes over time. We measured the extent of temporal change in each parasite community by calculating the cumulative distance moved by the parasite community in multivariate space. More specifically, for each sequential pair of surveys, we measured the Euclidean distance between the states (i.e., NMDS coordinates) of each community at those two times. The greater the distance between community states at two time points, the more the community has changed over that time interval. As a measure of cumulative temporal change, we then calculated the total trajectory length (De Cáceres et al. 2019), which is the sum of all distances between sequential community states over the entire sequence of field surveys. We tested whether symbiont exposure history impacted cumulative temporal change in the parasite community using an ANOVA that included the three symbiont inoculation treatments and all interactions between inoculations in our model. Cumulative temporal change was log10 transformed to minimize heteroscedasticity. The ANOVA were implemented in the ‘car’ package with type 3 tests. As a measure of effect size for ANOVAs we reported η^2^_partial_, which is the proportion of variance explained after accounting for the effects of other variables in the statistical model. Differences between treatment groups in the ANOVA were evaluated based on Tukey’s HSD.

### Parasite community divergence

When communities start in different states, and then are exposed to a common pool of propagules, they may converge in community structure over time. Thus, parasite communities may converge over time because, after receiving different initial symbiont inoculation treatments, all hosts were exposed to a common pool of parasite propagules for several weeks. To detect signals of community convergence, we first calculated the NMDS centroid of each treatment group in each survey, as a measure of the central tendency of the parasite communities. Second, for each pair of treatment groups, we measured the Euclidean distance between their community centroids at each survey time. Third, we analyzed the relationship of the distance between treatment centroids (i.e., dissimilarity in community states) and time, to evaluate whether community trajectories of the different treatment groups generally became more similar as the communities assembled (De Cáceres et al. 2019). To test for convergence or divergence between each pair of treatment groups, we used the non-parametric Mann-Kendall test (R package ‘trend’). This test detects monotonic trends in the relationship between dissimilarity (i.e., measured as distance between treatment centroids) and time. In this test, trajectories that are diverging over time have a tau value greater than 0 (i.e., positive relationship between dissimilarity and time), while trajectories that are converging over time have tau less than 0 (i.e., negative relationship between dissimilarity and time). Tau values are derived from the sign (positive or negative) of the slopes that describe community divergence; these values do not consider the magnitude of the slopes. To determine the magnitude of trends in community divergence, we also report the sens slope (function ‘sens.slope’ in the trend package), which is the median of the slope values. Note that the p-values from the tau-value trend tests also apply to the sens slopes.

### Drift

Finally, we tested whether there was evidence for drift among parasite communities and how the importance of drift changed during the experiment. Specifically, we assessed whether within-group variation increased over time by calculating each parasite community’s distance to the group centroid at the time of each survey. An increase in the distance from the centroid over time would indicate that these communities are diverging from a central community state. Such within-group divergence may chiefly represent ecological drift, as our experimental approach sought to eliminate within-group variation in other deterministic factors. We tested for drift among parasite communities using linear mixed effect models. We used plant ID as a random effect (random intercept only) to account for repeated measurements, and we log10 +1 transformed the distance to the centroid to minimize heteroscedasticity. To test our hypothesis that the importance of drift would decrease over time, we used a piecewise regression model to fit two mixed effect models: one fitting an earlier portion of the community trajectories, and one fitting a later portion. The piecewise approach does this by estimating the break point between the earlier and later mixed models as well as the parameters of each model, with each model’s slope being of primary interest (i.e., a change in slope would represent a change in drift). Mixed models were built in ‘nlme’ package, and the piecewise regression was conducted with the ‘segmented’ package. All analyses and graphics were produced in R v. 4.0.2 (R Core Team, 2021).

## Results

### Multivariate patterns of parasite community structure

Parasite community structure differed between symbiont inoculation treatments and over time (Figure 1, Figure S2). While all inoculation treatment effects and their interactions were statistically significant (PERMANOVA, p < 0.001), they explained little variation in parasite community structure among hosts (< 3% in total) (Table S4, S5). For example, the interaction between *Epichloë* and *Rhizoctonia* inoculations had the largest effect on parasite communities but still explained very little variation (*Epichloë* * *Rhizoctonia:* p < 0.001, R^2^ = 0.006). On the other hand, sampling day explained 8.6% of the variation in parasite communities (Table S4). Multivariate homogeneity of variance was not equal across sampling days (ANOVA, F12,1583 =19.32, p < 0.0001, Figure S3), nor among inoculation treatment groups (F7,1588 =5.06, p < 0.0001), so within-group variation across time and inoculation treatments were also important factors driving parasite community patterns. Among inoculation treatment groups, differences in multivariate variance were primarily driven by the hosts inoculated with only *Colletotrichum*, which had lower variation in parasite community structure relative to all other symbiont-inoculated hosts (post-hoc comparisons; Table S6).

**Figure 1.**
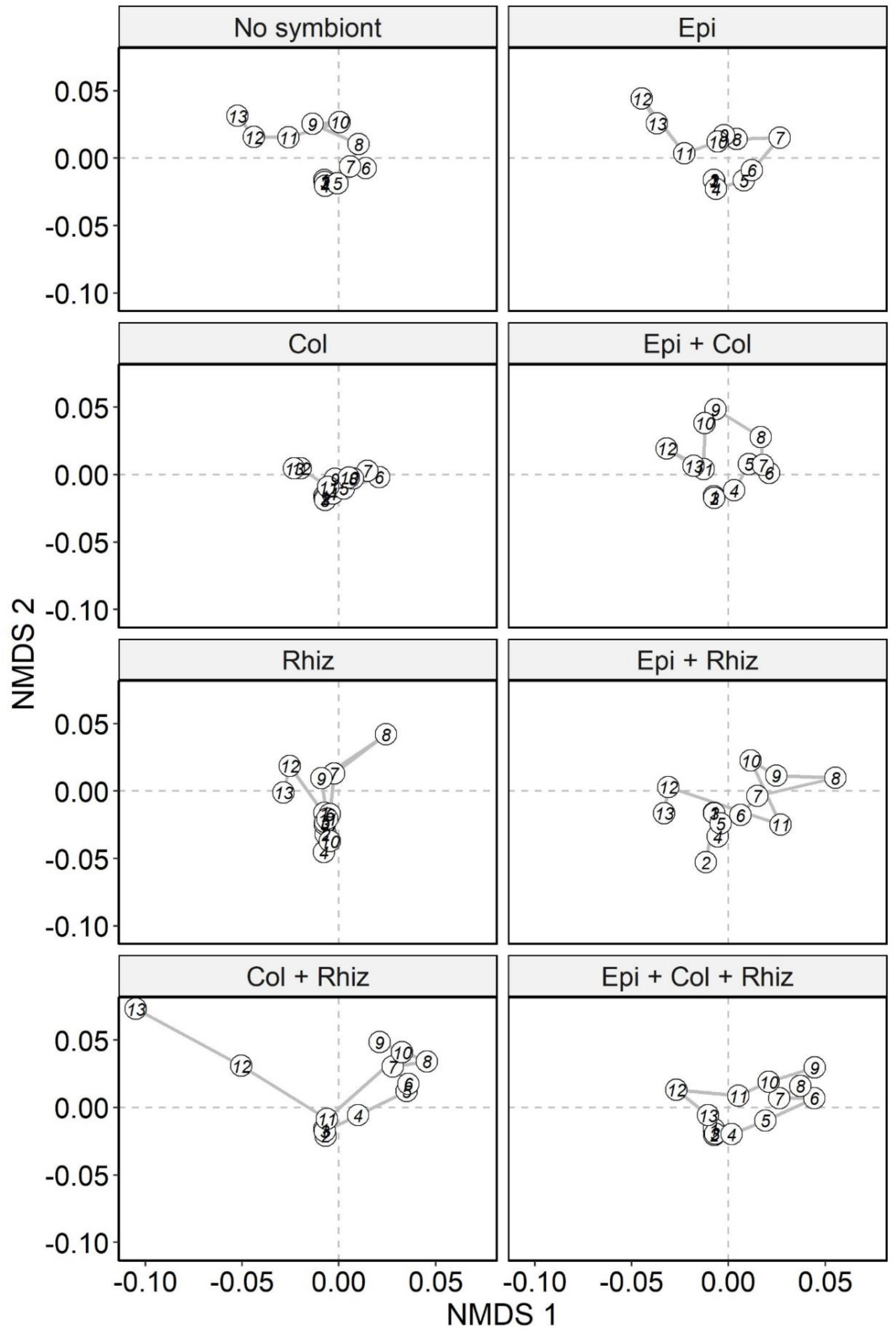
The trajectory of parasite community assembly within hosts inoculated with a factorial combination of three symbionts. Each panel shows parasite community patterns, based on an NMDS of Bray-Curtis distances, for an inoculation treatment group. Within these NMDS diagrams: *Colletotrichum* was associated with positive values of NMDS axis 1 and 2, *Rhizoctonia* was associated with negative values of NMDS axis 2, *Puccinia* was associated with negative values NMDS axis 1 and positive values of NMDS axis 2, and *Pyricularia* was associated with positive values of NMDS axis 1 and negative values of NMDS axis 2 (Table S2). Each circle represents the centroid of the parasite communities in that treatment group in one survey. Numbers within circles designate time (the sequence of surveys) and lines connect sequential surveys, showing the community trajectories.

### Temporal change in parasite community structure

The cumulative temporal change of parasite communities was decreased by *Colletotrichum* inoculation (ANOVA, F_1,125_ = 5.88 p = 0.017, η^2^_partial_ effect size = 0.045), and there was an interaction between *Colletotrichum* and *Rhizoctonia* inoculations (F_1,125_ = 4.74, p = 0.031, η^2^_partial_ = 0.037), where in plants inoculated with *Colletotrichum*, co-inoculation with *Rhizoctonia* increased the temporal change in parasite community structure (Figure 2). There was also a marginally significant three-way interaction wherein inoculation with all three symbionts tended to decrease temporal change in parasite communities (F_1,125_ = 2.65, p = 0.106, η^2^_partial_ = 0.021). There was no main effect of *Rhizoctonia* or *Epichloë* inoculations on the temporal change of parasite communities, and all other interactions among symbiont inoculations were non-significant (p > 0.05; Table S7). Pairwise comparisons indicate plants inoculated with only *Colletotrichum* harbored parasite communities that changed less over the course of the entire experiment relative to plants inoculated with both fungal parasites, *Colletotrichum* and *Rhizoctonia* (Tukey HSD, p = 0.02), but not when these parasites were in coinfection with *Epichloë* (Figure 2). Overall, these results suggest that initial exposure to multiple parasites has the potential to drive greater long-term community change, but only in limited contexts determined by exposure to other symbionts.

**Figure 2.**
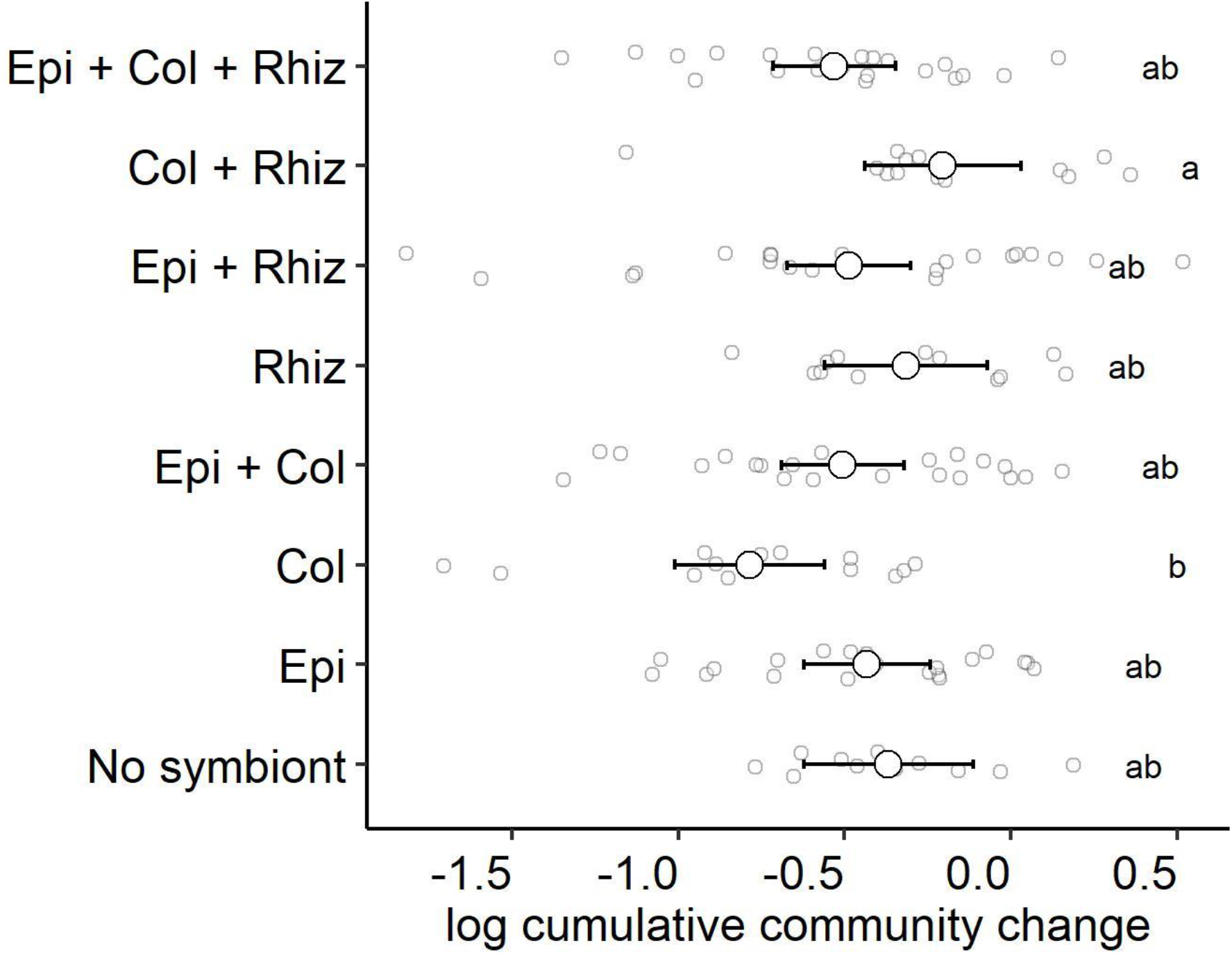
The cumulative temporal change of parasite communities (i.e., the total Euclidean distance moved by a parasite community) differed between inoculation treatment groups. Post hoc comparisons based on Tukey HSD (grouping denoted by letters) indicate that plants co-inoculated with *Colletotrichum* and *Rhizoctonia* harbored parasite communities that changed more when compared to *Colletotrichum* inoculated plants. Plotted are the treatment means and their 95% confidence intervals. Smaller points display the raw data.

**Figure 3.**
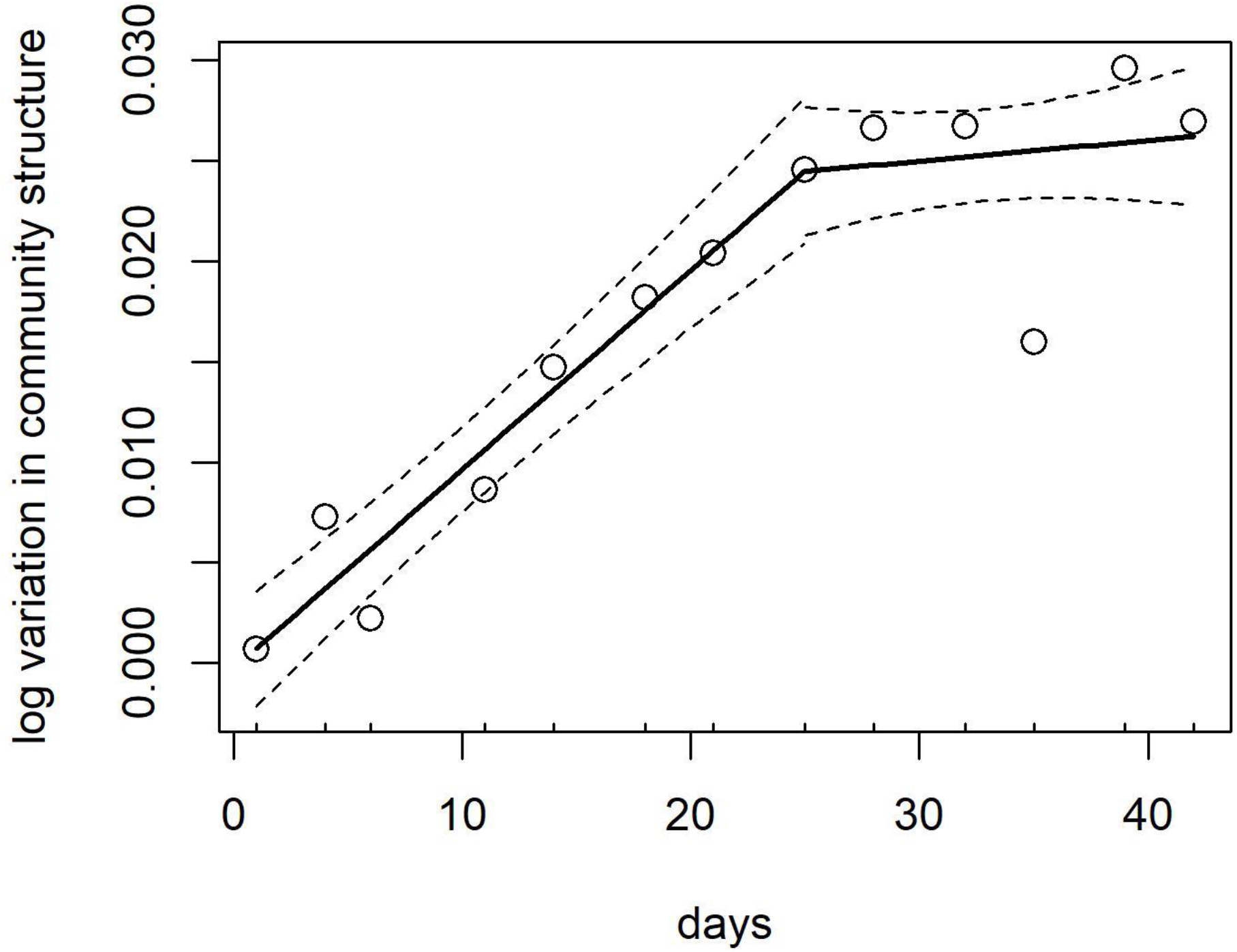
Early in the experiment, the variation in parasite community structure among hosts increased over time, consistent with ecological drift. The increase in variation of community structure was no longer apparent after day 25 of the field survey, consistent with the prediction that drift would be more important earlier in the experiment. Plotted is the fitted line (solid) and 95% confidence interval (dashed lines) from the piecewise regression mixed model, and each point is the mean variation in community structure across all hosts (i.e., log transformed distance to the centroid) at a given survey event.

### Parasite community divergence

Despite exposing hosts to a common pool of parasite propagules in the field, which we expected to generate parasite community convergence, trend tests indicated that parasite community trajectories generally diverged between symbiont inoculation treatment groups. Notably, no treatment groups showed any sign of convergence, as no values of tau were less than zero (tau range: 0.1 to 0.92, Figure S4-A). These results indicate persistent changes in parasite community structure over time in response to their initial symbiont inoculations. Moreover, plants that were not inoculated with any symbiont exhibited parasite community trajectories that significantly diverged from those of plants in all other symbiont inoculation treatments (mean tau: 0.56, range: 0.36 to 0.87, p <0.05) except one (*Colletotrichum* and *Rhizoctonia*, tau = 0.36, p value =0.10, Figure S5). Further, inoculation with only *Epichloë*, the hypothesized mutualist, yielded parasite communities that significantly diverged from all other inoculation treatments (Tau range: 0.46 to 0.92, p< 0.05). On the other hand, in the absence of *Epichloë*, parasite community trajectories from plants inoculated with only the parasite *Rhizoctonia* did not show significant trends of divergence from the other parasite inoculation treatments (i.e., co-inoculated with *Colletotrichum* and inoculated with only *Colletotrichum*) (p > 0.05).

The magnitude of parasite community divergence, measured by the sens slope, also varied among treatment groups. Notably, plants co-inoculated with both parasite species, *Colletotrichum* and *Rhizoctonia*, but not with *Epichloë*, had the greatest mean sens slope value (0.004), indicating that parasite communities in that treatment group diverged the most, on average, from the other treatment groups (Figure S4-B). In particular, the greatest rate of divergence (sens slope = 0.0057) was between parasite communities from plants co-inoculated with both parasites, *Rhizoctonia* and *Colletotrichum*, and parasite communities from plants inoculated with only *Colletotrichum* (p < 0.001). The sign of the sens slope is linked to Mann-Kendall trend test, so just as for the tau values, no sens slope values were less than zero, again indicating no convergence. Overall, both the sens slope and trend test indicated that parasite community trajectories diverged in response to a host’s initial exposure to symbionts. Moreover, the magnitude of divergence depended on the symbiont species inoculated, as we did not observe significant effects of *Rhizoctonia* inoculation alone on parasite community patterns and the magnitude of divergence was greatest in hosts co-inoculated with *Rhizoctonia* and *Colletotrichum*.

### Drift

Ecological drift was also an important driver of variation in parasite community structure, but only in the beginning of the experiment. Initially, the distance of the parasite communities to their treatment-group centroid increased over time (slope = 0.0010, 95% CI = (0.0008, 0.0012)), indicating increased variation in parasite community structure within treatment groups over time. However, piecewise regression identified a break point in our drift model, where after day 25 of the field survey (break point estimate = day 25, 95% CI = (20.1, 29.8)), the distance from the centroid no longer increased (slope = 0.0001, 95% CI = (−0.0002, 0.0004)) (Figure 4). This finding provides evidence of ecological drift, and supports the prediction that drift is more important earlier in community assembly.

## Discussion

Community ecology theory posits that variation in community structure is driven by four core processes (Vellend 2010), and we aimed to explore the role of dispersal (i.e., order of infection), selection (i.e., species interactions), and ecological drift (i.e., among host divergence), in structuring parasite communities, as the roles of the four processes are rarely all considered simultaneously. Using experimental inoculations and longitudinal field infection data, we showed that prior exposure of hosts to a combination of symbionts alters parasite community composition patterns. Importantly, plants that were not inoculated with symbionts or only inoculated with the mutualist had divergent parasite community trajectories relative to other symbiont-inoculated plants, suggesting selection is contributing to parasite community assembly. Along with these deterministic processes, parasite communities also varied considerably among hosts over time, which is indicative of drift. As predicted by theory, evidence of drift was only apparent in the beginning of the experiment. Taken together, these data provide evidence of persistent changes in parasite community structure based on initial variation in parasite community composition, and selection along with drift are important for parasite diversity.

### Host exposure to symbionts alters parasite community assembly

The initial composition of symbionts within a host can generate priority effects that alter a host’s susceptibility to future infection (Graham 2008, Ezenwa et al. 2010, Syller and Grupa 2016, Halliday et al. 2018). Within our system, priority effects have been described to alter parasite transmission among hosts in the field (Halliday et al. 2017) and parasite growth rates within host individuals in laboratory settings (O’Keeffe et al. 2021). At the community-level, differences in infections due to within-host priority effects may also result in divergent parasite community trajectories (Benesh and Kalbe 2016, Halliday et al. 2020). Consequently, our primary goal was to detect trends of convergence or divergence in parasite community patterns when hosts are initially exposed to a combination of symbionts. In our experiment, hosts were exposed to a common pool of parasites in the field after inoculations and should experience similar rates of transmission, so communities may converge. Yet, parasite communities from symbiont inoculated hosts tended to diverge from each other, and there was no signal of convergence between communities. Thus, our results show that exposing hosts to certain combination of symbionts is often enough to generate differences in parasite communities that persisted over time.

While we generally observed divergence between parasite community trajectories, plants inoculated with only one of the parasites, *Rhizoctonia*, showed no clear signal of divergence when compared to the other parasite inoculated hosts. Further, the magnitude of divergence (i.e., the sens slope) was lowest for this treatment group. When no strong signals of divergence were found, parasite communities diverged in the beginning and then converged and diverged again, resulting in no consistent signal (i.e., positive and negative slopes). Consequently, at different time points we may detect differences or similarities between parasite communities, but longer-term sampling (over 6 weeks, the lifespan of leaves/hosts) shows that these directional changes were not consistent. We hypothesize the observed non-directional changes in *Rhizoctonia* inoculated hosts are driven by the necrotrophic feeding strategy of this parasite. Plants inoculated with *Rhizoctonia* experience a higher rate of leaf mortality (O’Keeffe et al. 2021). The high turnover of leaf hosts may lead to less consistent community patterns, because parasite communities are more frequently going extinct and restarting community assembly. We interpret this as prior infections by *Rhizoctonia* lowering community stability through increases in host mortality and resulting in greater temporal change in parasite community structure. Overall, these dynamics can manifest as transient community states that are related to priority effects (Fukami 2015).

### Symbiont coinfections

Parasite communities from hosts initially co-inoculated with both parasites became infected with parasite communities that changed more over time, and generally these communities diverged at a faster rate. In particular, the greatest differences in parasite community patterns occurred between hosts co-inoculated with both parasites and *Colletotrichum-* only inoculated hosts. At one end, inoculation with only *Colletotrichum* resulted in parasite communities that did not vary much through time, but when exposed to both parasite species, the resulting parasite communities changed more. Coinfection with *Rhizoctonia* and *Colletotrichum* has been shown to affect the outcome of parasite interactions. Specifically, field studies demonstrated an antagonistic relationship between these parasites, hypothesized to occur through resource competition (Halliday et al. 2017, 2018), while later laboratory studies provided evidence for facilitation of *Rhizoctonia* by *Colletotrichum* when in coinfection (O’Keeffe et al. 2021). Along with these changes in species interactions, our study shows that initial coinfection by these parasites also translate to differences in parasite communities over time.

Parasite community patterns and some symbiont inoculation treatment effects were altered when in the presence of the mutualist *Epichloë*. Specifically, the increase in temporal change in parasite communities due to parasite co-inoculation was not apparent in *Epichloë* inoculated hosts. At the same time, parasite community trajectories from *Epichloë* hosts diverged from all other symbiont inoculations. The prevalence of *Epichloë* in the natural host population is high (~ 85%), and thus these results are relevant for hosts in the field. However, the high prevalence of *Epichloë*, combined with its potential to diminish some effects of parasite inoculation, may make it difficult to detect some of the effects observed in these inoculation experiments in naturally occurring host populations. Nonetheless, considering the effects of other symbionts is important because the outcome between parasite can be modified in the presence of mutualist symbionts (Marchetto and Power 2018, O’Keeffe et al. 2021). Within our system, the two parasites and *Epichloë* stimulate phytohormone pathways that alter parasite establishment and growth (Saikkonen et al. 2010). For example, exposure to salicylic acid, which is hypothesized to increase in response to endophyte infections (Saikkonen et al. 2013), modified the outcome of interactions between parasites in tall fescue by decreasing coinfections and increasing disease severity (Halliday et al. 2018). These data suggest that immune mediated interactions that may occur when in coinfection with a mutualist may be of broader importance for parasite community patterns.

### Drift contributes to parasite diversity during community assembly

Among host individuals, the distance from the parasite community centroid initially increased over time, indicative of the importance of drift during the early phases of infection Given the high degree of aggregation commonly reported across and within parasite species (Poulin 2007), it is not surprising that most parasite communities exhibit a high degree of among host divergence, because most hosts have low parasite population sizes. Parasite aggregation is largely a product of host heterogeneity i.e., variation in host exposure, behavior, and genetics (Johnson and Hoverman 2014). We attempted to minimize these sources of host heterogeneity in our study by continually moving plants within the field to homogenize parasite exposure and by using plants from the same cultivar to reduce genetic diversity. Although we attempted to reduce sources of among-host variation, we still detected a large amount variation in parasite community structure among hosts, which are likely due to stochastic colonization and extinction of parasite species.

The role of drift in structuring parasite communities has received little attention in disease ecology (Seabloom et al. 2015), and some evidence suggests drift is not important for parasites with complex lifecycles (Moss et al. 2020). Nevertheless, drift has the potential to interact with the other relevant drivers of parasite diversity. For example, if selection is weak during the early phases on infection, potentially due to low population sizes or overall weak interaction strength, drift may be a chief driver of parasite diversity patterns (Vellend 2010). These relationships are likely dynamic through time, and as parasite population sizes and interaction strength increase more deterministic processes (e.g., selection) will contribute to parasite diversity and drift becomes less important. Ultimately, we predict the contribution of drift is an important factor contributing to parasite diversity and the role of stochastic processes should further be considered in the context of parasite community assembly.

While symbiont exposure history had clear effects on parasite community assembly, our results may vary depending on the timing and environmental context of our experiment. Throughout the growing season hosts are exposed to different subsets of parasite species that vary in dispersal rates and timing. In our system, parasite phenology varies across the three main foliar fungal parasites within this population of tall fescue (Halliday et al. 2017). We conducted our experiment during the early fall to ensure all three parasites were readily present in the host population and could infect our host plants. Since the general pool of parasite and their transmission vary over time (Halliday et al. 2017), it is likely our results are contingent on time of year we deployed our lab inoculated plants into the field. Further, differences in parasite phenology coincide with differences in precipitation and temperature that may modify the strength of parasite interactions and alter community patterns. Given that both the environmental context and disease pressure are dynamic throughout time, these results are likely sensitive to the timing of our experiment.

### Conclusion

Initial variation in parasite community composition due to dispersal can drive selection and subsequently parasite community assembly (Benesh and Kalbe 2016, Halliday et al. 2020). In our experiment, a host’s initial inoculum of symbionts was important for parasite diversity in the field. In particular, co-inoculation with both parasite species were important drivers of parasite community change. While our experiment focused on the role of selection (i.e., symbiont inoculations), parasite communities also tended to diverge from each other in the beginning of the experiment due to drift., drift may interact with other processes that shape parasite diversity patterns and merits further consideration. Overall, a better understanding of these factors as structuring forces of parasite communities can contribute to our understanding of disease outcomes in natural populations, because variation in parasite community structure can have implications for host health and disease spread (Jolles et al. 2008, Hoverman et al. 2013).

## Acknowledgments

We thank Dr. Tim Phillips and particularly Dr. Rebecca McCulley for generously providing tall fescue seed. We also thank the Mitchell lab, Peter Morin, Cara Faillace, and members of the Morin lab for their feedback on this manuscript. This work was supported by the NSF-USDA joint program in Ecology and Evolution of Infectious Diseases (USDA-NIFA AFRI grant 2016-67013-25762).

## Author contributions

R.L.G. analyzed the data and wrote the manuscript. B.N.J. performed the experiment. C.E.M. conceptualized and designed the experiment. All authors contributed to revising the manuscript.

