## Supplemental Material for "Host exposure to symbionts and ecological drift generate divergence in parasite community assembly"

**Supporting information for "Host exposure to symbionts and ecological drift generate divergence in parasite community assembly"**

Rita L. Grunberg, Brooklynn N. Joyner, and Charles E. Mitchell

**Appendix I. Methods for symbiont inoculations**

1. **Endophyte inoculation:** The *Epichloë*-free treatment group comprised 52 plants grown from seeds produced by Tim Phillips at University of Kentucky, and the *Epichloë* inoculation group comprised 84 plants (a larger number because we anticipated <100% transmission of the endophyte from seed) grown from seeds produced by the Noble Research Institute in Oklahoma (Table S1). All seeds used in the experiment were of the cultivar KY-31.
2. **Parasite inoculations:** Our sources of inoculum for each of the parasite species were derived from cultures collected in Duke Forest in 2015 by F. Halliday and K. O'Keeffe. For the *R. solani* inoculation, we placed a mycelium PDA plug at the base of the oldest living leaf to establish infection. Plants that were not inoculated with *R. solani* were mock-inoculated with a blank PDA plug. The inoculation site was then covered with moist cotton and wrapped in aluminum foil and parafilm to maintain a humid environment for parasite growth. After two days, we removed the inoculum covering. *Colletotrichum cereale* inoculations were implemented according to Beirn et al., 2015. We established C. *cereale* PDA cultures with mycelium plugs that were grown out on plates for 10 days. We then flooded and scraped these plates to produce a mycelial and spore solution. This growth and flooding process was repeated until we reached a spore density of $10^6$ conidiospores/mL water. Plants were then inoculated with C. *cereale* by brushing 10 mL of the spore solution onto the oldest leaf. Plants that were not inoculated with the *C. cereale* were mock-inoculated with a solution of autoclaved water and potato dextrose broth. We then maintained plants of all treatments for two days in dew chambers, which consisted of a misted sealed bag kept over each individual plant.

3. **Observation of symptoms on inoculated leaves:** To assess symptoms due to parasite inoculations, we tested whether the symbiont inoculations affected the lifespan (i.e., number of days surveyed in the field) of the inoculated leaf using an ANOVA. We included all symbiont inoculations and their interactions in the ANOVA. The parasite inoculations negatively affected the lifespan of the lab-inoculated leaves (Figure S1). We detected a main effect of inoculation by both parasites, *Rhizoctonia (*ANOVA, $F_{1,125} = 5.16$, $p = 0.025$, $\eta^2_{partial} = 0.039$) and *Colletotrichum* ($F_{1,125} = 7.89$, $p = 0.006$, $\eta^2_{partial} = 0.061$) on inoculated leaf lifespan. Further, there was an interaction between both parasites, with co-inoculations having the greatest negative effect on leaf lifespan (*Rhizoctonia* * *Colletotrichum*, $F_{1,125} = 45.63$, $p = 0.0002$, $\eta^2_{partial} = 0.105$). There was no main effect of *Epichloë* on leaf lifespan ($F_{1,125} = 0.60$, $p = 0.44$, $\eta^2_{partial} = 0.003$), and all other interactions between *Epichloë* and the parasite inoculations were not significant ($p > 0.05$) (Table S2). Together, leaf inoculation treatments explained 22% of the variation in inoculated leaf lifespan.

### Appendix II: Additional description of plot design

Lab inoculated plants were maintain in individual pots and outplanted into 1.5 x 12m plot at Weidner Farm. This plot was fenced to exclude deer, with an additional mesh barrier to exclude small mammals. Within the plot, there were three rows of 47 holes each and these holes were arranged 0.25 m apart along a 12m length of the plot. Plants were placed into holes randomly within the plot and each plant was relocated to a new hole every week to homogenize parasite exposure. The plot contained a dense cover of resident tall fescue (~ 80% cover within the plot) to act as a source of parasite inoculum for the experimental plants.

**Table S1.** Sample sizes of experimentally inoculated plants. Here, N (initial) is the total plants used for each inoculation treatment group, N (analyzed) indicates total number of plants that showed disease symptoms during the field survey and retained for statistical analyses, N (*Epichloë*) indicates the total number of plants that tested positive for *Epichloë* at the end of the experiment, and % infected indicate the percentage of symptomatic plants that tested positive for Epichloe.

| **Inoculation treatment** | **N (**initial**)** | **N (**analyzed**)** | **N (***Epichloë* **)** | **% infected (***Epichloë***)** |
|---|---|---|---|---|
| No symbiont | 12 | 11 | 0 | 0 |
| Col | 14 | 14 | 0 | 0 |
| Col + Rhiz | 13 | 13 | 0 | 0 |
| Rhiz | 13 | 12 | 0 | 0 |
| Epi | 21 | 20 | 9 | 45 |
| Epi + Col | 21 | 21 | 5 | 24 |
| Epi + Col + Rhiz | 21 | 21 | 5 | 24 |
| Epi + Rhiz | 21 | 21 | 1 | 5 |

**Table S2.** ANOVA table evaluating differences in leaf lifespan among symbiont inoculation treatments. $\eta^2_{partial}$ is reported as an effect size, and bolded p-values are statistically significant ($p < 0.05$).

| | DF | F | p-value | $\eta^2_{partial}$ |
|---|---|---|---|---|
| Epi | 1 | 0.596 | 0.442 | 0.003 |
| Col | 1 | 7.892 | **0.006** | 0.061 |
| Rhiz | 1 | 5.158 | **0.025** | 0.039 |
| Epi*Col | 1 | 0.357 | 0.551 | 0.003 |
| Epi*Rhiz | 1 | 0.329 | 0.567 | 0.002 |
| Col*Rhiz | 1 | 14.736 | **0.0002** | 0.105 |
| Epi*Col*Rhiz | 1 | 1.049 | 0.308 | 0.008 |

**Table S3.** Parasite species scores (including the dummy variable) association with NMDS axes. Note that species are also plotted in the NMDS diagram in Figure S2.

| Parasite | NMDS1 | NMDS2 |
| --- | --- | --- |
| *Colletotrichum cereale* | 0.3858 | 0.1938 |
| *Rhizoctonia solani* | -0.0549 | -0.5193 |
| *Puccinia coronata* | -0.3637 | 0.3160 |
| *Pyricularia grisea* | 0.5252 | -0.8485 |
| Dummy | -0.0022 | -0.0026 |

**Table S4.** Summary table of PERMANOVA results based on Bray-Curtis distances evaluating the effects of symbiont inoculations and survey event on parasite communities infecting tall fescue.

| Inoculation treatment | Df | Pseudo F | $R^2$ | P-value |
|---:|---|---|---|---|
| Epi | 1 | 1.0223 | 0.00053 | 0.001 |
| Col | 1 | 4.8094 | 0.00252 | 0.001 |
| Rhiz | 1 | 4.4138 | 0.00231 | 0.001 |
| Survey event | 12 | 13.8094 | 0.0867 | 0.001 |
| Epi * Col | 1 | 0.1539 | 0.00008 | 0.001 |
| Epi * Rhiz | 1 | 10.9661 | 0.00574 | 0.001 |
| Col * Rhiz | 1 | 1.4486 | 0.00076 | 0.001 |
| Epi * Col * Rhiz | 1 | 4.2232 | 0.00221 | 0.001 |

**Table S5.** Summary table of pairwise PERMANOVAs conducted between each inoculation treatment group. Bolded are the p-values that are statistically significant p < 0.05, and the listed p-values were adjusted for multiple comparisons.

| | No symbiont | Col | Col + Rhiz | Rhiz | Epi | Epi Col | Epi + Col + Rhiz | Epi + Rhiz |
|---|---|---|---|---|---|---|---|---|
| No symbiont | - | - | - | - | - | - | - | - |
| Col | **0.025** | - | - | - | - | - | - | - |
| Col + Rhiz | **0.033** | **0.025** | - | - | - | - | - | - |
| Rhiz | 0.280 | 0.432 | **0.025** | - | - | - | - | - |
| Epi | 0.557 | **0.028** | **0.037** | **0.037** | - | - | - | - |
| Epi + Col | 0.186 | **0.028** | 0.556 | 0.054 | 0.354 | - | - | - |
| Epi + Col + Rhiz | **0.025** | **0.025** | 0.061 | **0.025** | **0.025** | **0.025** | - | - |
| Epi + Rhiz | **0.044** | 0.587 | **0.028** | 0.401 | **0.034** | **0.037** | 0.069 | - |

**Table S6.** Tukey post-hoc comparisons for test of multivariate homogeneity of variance in relation to the PERMANOVA. Comparisons are conducted between inoculation treatment groups. P-values are adjusted for multiple comparisons and bolded values are significant ($p < 0.05$).

| **Comparison** | **p-value** |
|---:|---:|
| Col V. no symbiont | 0.195 |
| Col + Rhiz V. no symbiont | 0.061 |
| Rhiz V. no symbiont | 0.992 |
| Epi V. no symbiont | 1.000 |
| Epi + Col V. no symbiont | 0.991 |
| Epi + Col +Rhiz V. no symbiont | 1.000 |
| Epi + Rhiz V. no symbiont | 0.901 |
| Col + Rhiz V. Col | **< 0.001** |
| Rhiz V. Col | **0.016** |
| Epi V. Col | **0.035** |
| Epi + Col V. Col | **0.004** |
| Epi + Col + Rhiz V. Col | **0.024** |
| Epi + Rhiz V. Col | **0.001** |
| Rhiz V. Col + Rhiz | 0.389 |
| Epi V. Col + Rhiz | **0.039** |
| Epi + Col V. Col + Rhiz | 0.167 |
| Epi + Col + Rhiz V. Col + Rhiz | 0.059 |
| Epi + Rhiz V. Col + Rhiz | 0.463 |
| Epi V. Epi + Rhiz | 0.998 |
| Epi + Col V. Rhiz | 1.000 |
| Epi + Col + Rhiz V. Rhiz | 0.999 |
| Epi + Rhiz V. Rhiz | 1.000 |
| Epi + Col V. Epi | 0.998 |
| Epi + Col + Rhiz V. Epi | 1.000 |
| Epi + Rhiz V. Epi | 0.929 |
| Epi + Col + Rhiz V. Epi + Col | 1.000 |
| Epi +Rhiz V. Epi + Col | 0.999 |
| Epi + Rhiz V. Epi + Col + Rhiz | 0.965 |

**Table S7.** ANOVA table evaluating differences in cumulative temporal change of parasite communities (measured as Euclidean distance moved by a parasite community) among symbiont inoculation treatments. $\eta^2_{partial}$ is reported as an effect size, and bolded p-values are statistically significant ($p < 0.05$).

| | ANOVA Type 3 Test | | | |
|---|---|---|---|---|
| | DF | F | p-value | $\eta^2_{partial}$ |
| Epi | 1 | 0.163 | 0.687 | 0.001 |
| Col | 1 | 5.880 | **0.017** | 0.045 |
| Rhiz | 1 | 0.087 | 0.769 | 0.001 |
| Epi*Col | 1 | 2.499 | 0.116 | 0.020 |
| Epi*Rhiz | 1 | 0.231 | 0.631 | 0.002 |
| Col*Rhiz | 1 | 4.743 | **0.031** | 0.037 |
| Epi*Col*Rhiz | 1 | 2.650 | 0.106 | 0.021 |

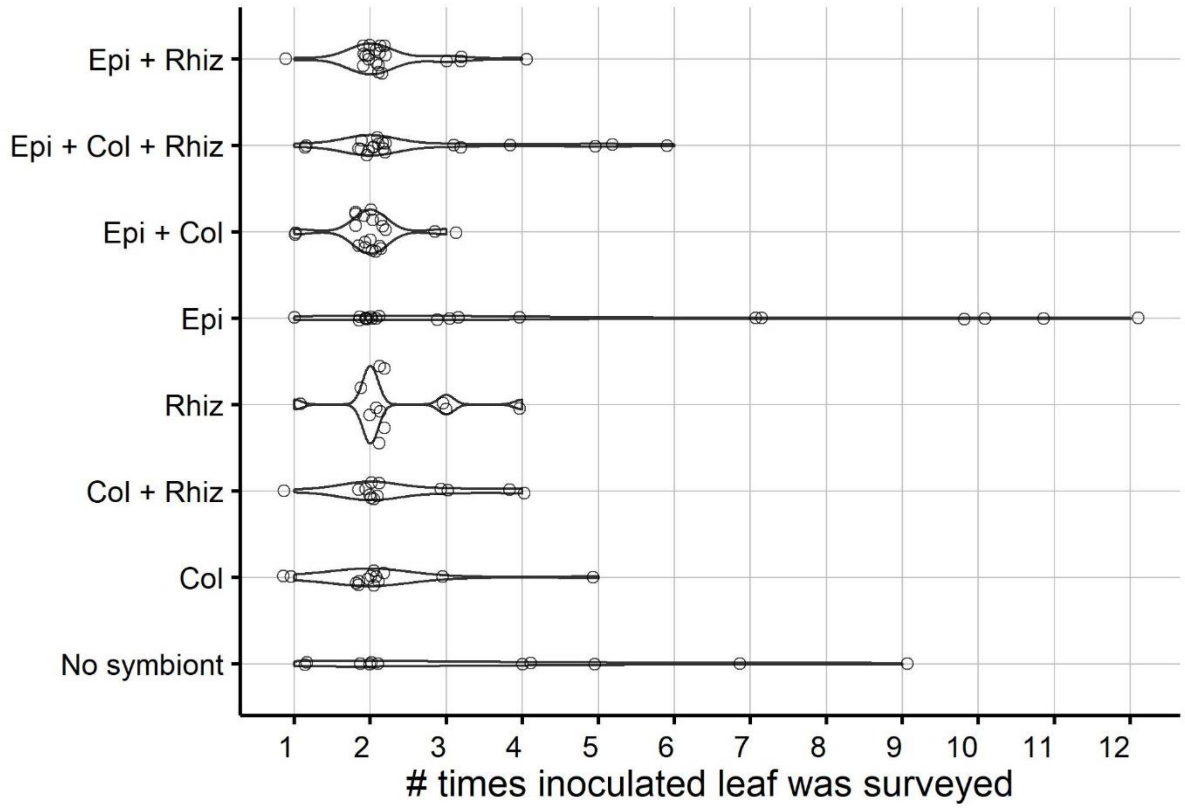


**Figure S1.** Parasite inoculations resulted in a decline in inoculated leaf lifespan (i.e., the numbers of times an inoculated leaf was surveyed). Each point indicates an individual leaf that received an inoculation treatment and how many times that leaf was surveyed in the field. Inoculations were implemented on the oldest living leaf prior to being outplanted in the field. Note that points are jittered to show the distribution of the raw data.

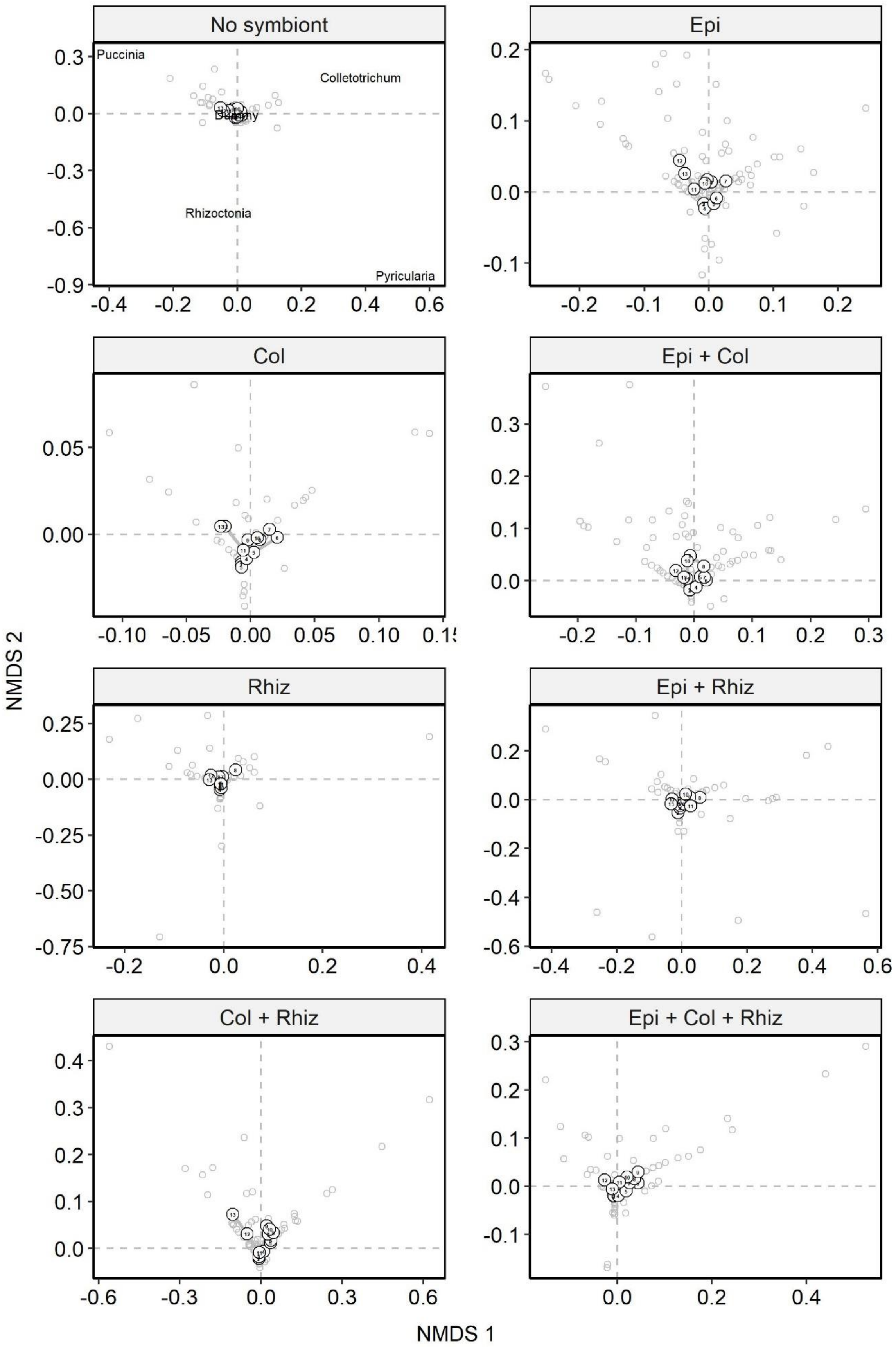
No symbiont
Puccinia
Colletotrichum
Rhizoctonia
Pyricularia
Epi
Col
Epi + Col
Rhiz
Epi + Rhiz
Col + Rhiz
Epi + Col + Rhiz
NMDS 2
NMDS 1

**Figure S2.** Parasite community patterns differed between symbiont inoculation treatments. Each panel depicts the NMDS diagram for an inoculation treatment group (stress= 0.04). Each large circle represents the centroid of the parasite communities in that treatment group in one survey, and smaller grey circles are the raw data from all hosts within a treatment group. Numbers within the larger circles designate time (the sequence of surveys) and lines connect sequential surveys, showing the community trajectories. The mapping of the parasite species scores (and dummy variable) are shown in the 'no symbiont' treatment panel. Note NMDS axes differ between inoculation treatment panels to show the spread of the raw data.

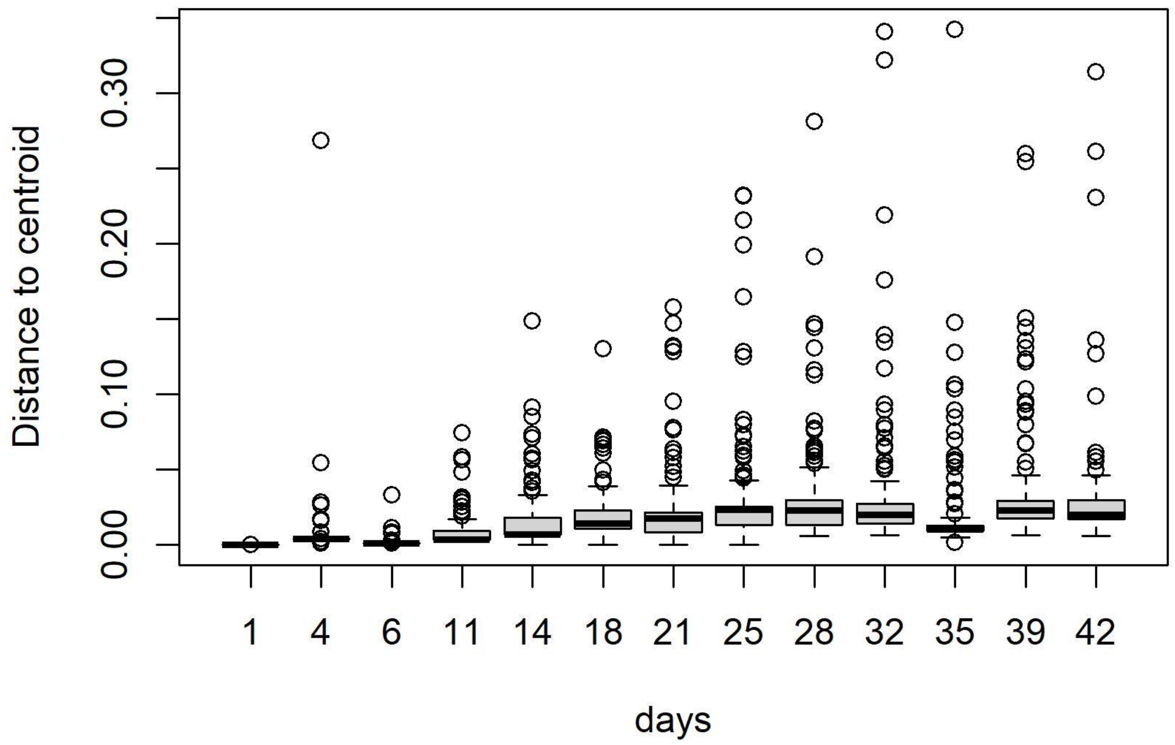


**Figure S3.** The distribution of distances to the parasite community centroid (shown is the raw data i.e., not log-transformed), displayed as boxplots, over time. These data were used to quantify variation in parasite communities over time in the drift model.

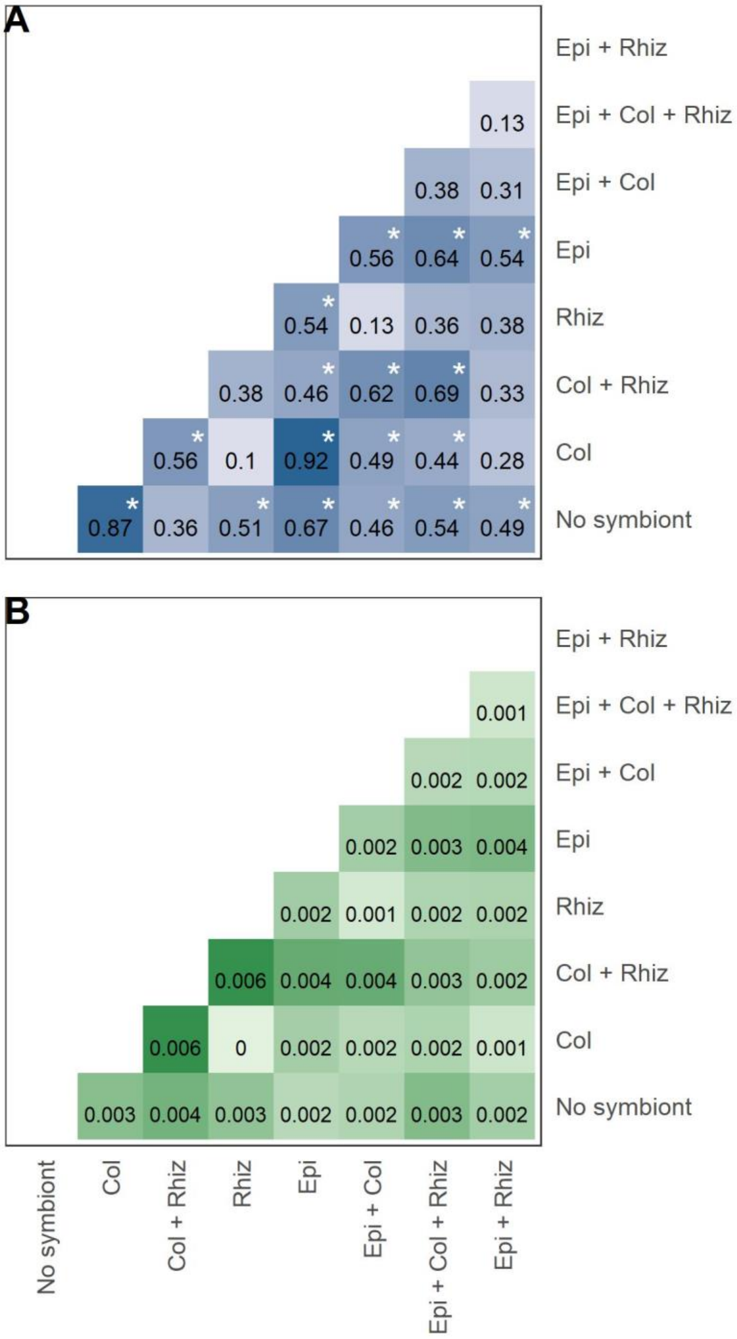


**Figure 4A-B.** Parasite community trajectories showed A) trends of divergence between most inoculation groups, and B) the magnitude of divergence between communities was greatest for parasite communities from hosts co-inoculated with both parasite species, *Colletotrichum* and *Rhizoctonia*. Panel A shows the results of pairwise Man-Kendell trend tests, which test for trends of convergence and divergence between communities. Within cells are Tau values, which indicate whether trajectories show signals of convergence (Tau < 0) or divergence (Tau > 0), and the fill of the cells correspond to Tau values, where white cells are 0 and darker blue cells are closer to 1. Panel B contains the values of pairwise sens slopes, which measures the magnitude of divergence. Within cells are the sens slope values, where darker green cells indicate a greater magnitude of divergence between community trajectories. Asterisks in panel A denote significant (p<0.05) trends of convergence or divergence, and these p-values also correspond to sens slope in panel B.

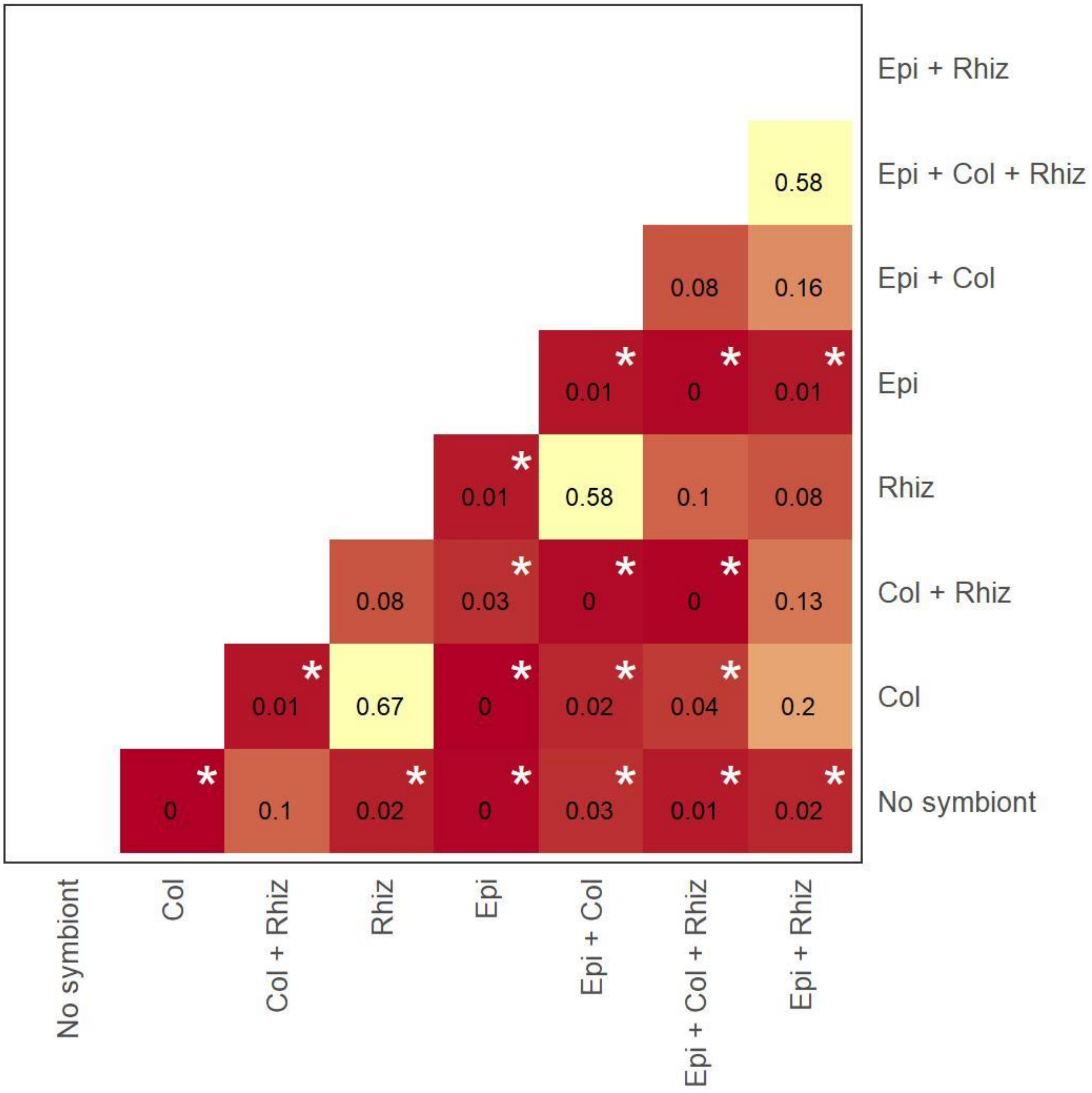


**Figure S5.** P-values associated with pairwise Man-Kendell trend tests and sens slope between parasite community trajectories for each treatment group. Asterisks denote significant ($p<0.05$) trends and sens slope. Within cells are p-values and the fill of the cells correspond to p-values, where yellow cells are 1 (non-significant trend) and darker red cells are closer to 0.
